# Metabolomics of aging in primary fibroblasts from small and large breed dogs

**DOI:** 10.1101/2021.02.25.432888

**Authors:** Paul S. Brookes, Ana G. Jimenez

## Abstract

Among several animal groups (eutherian mammals, birds, reptiles) lifespan positively correlates with body mass over several orders of magnitude. Contradicting this pattern are domesticated dogs, with small dog breeds exhibiting significantly longer lifespans than large dog breeds. The underlying mechanisms of differing aging rates across body masses are unclear, but it is generally agreed that metabolism is a significant regulator of the aging process. Herein, we performed a targeted metabolomics analysis on primary fibroblasts isolated from small and large breed young and old dogs. Regardless of size, older dogs exhibited lower glutathione and ATP, consistent with a role for oxidative stress and bioenergetic decline in aging. Furthermore, several size-specific metabolic patterns were observed with aging, including: (i) An apparent defect in the lower half of glycolysis in large old dogs at the level of pyruvate kinase. (ii) Increased glutamine anaplerosis into the TCA cycle in large old dogs. (iii) A potential defect in co-enzyme A biosynthesis in large old dogs. (iv) Low nucleotide levels in small young dogs that corrected with age. (v) An age dependent increase in carnitine in small dogs that was absent in large dogs. Overall, these data support the hypothesis that alterations in metabolism may underlie the different lifespans of small versus large breed dogs, and further work in this area may afford potential therapeutic strategies to improve the lifespan of large dogs.

## Introduction

Domestic dogs are the most morphologically and physiologically diverse mammal species known (Jimenez and Downs 2020), and the underlying consequences of this diversity span all levels of organization. Small breed dogs live significantly longer than large breed dogs (Jimenez 2016), and at the cell level this whole-animal trait is accompanied by increased cellular aerobic metabolism and mitochondrial proton leak as dogs of each size age, with increases in glycolytic rate in large breed dogs across their life span (Jimenez et al., 2018). Furthermore, small breed dogs accumulate more circulating lipid oxidative damage compared with large breed dogs, a pattern that would appear to oppose increased lifespan, although not uncommon in the animal kingdom (e.g., naked mole rats and bats also live long lives with accumulated damage (Jimenez and Downs, 2020)). While some progress has been made in elucidating potential mechanisms that dictate aging rates in domestic dogs, more work is needed in this area.

The metabolome is defined as the collection of metabolites in a cell or organism (Mishur and Rea 2012). To better understand the complexity of changes during aging, a metabolome-wide approach can be applied to pinpoint key metabolic steps that may determine lifespan in dogs (Soltow et al., 2010; Tombline et al., 2019; Hoffman et al., 2020). In rats, the plasma metabolome associated with aging showed changes in amino acids and a decline in lipid metabolism with increasing age, while in urine several Krebs’ cycle metabolites decreased with age (Mishur & Rea 2012). In dogs, metabolomic analysis has mainly been at the level of urine and plasma, and includes a life-long project on Labrador retrievers stratified as control animals versus those calorically restricted (CR) for life. Creatine increases were linked to muscle wasting with increased age (Wang et al., 2007) and lactate concentration in urine increased in older dogs (Wang et al. 2007), demonstrating a shift into a glycolytic phenotype similar to our studies using primary fibroblast cells (Jimenez et al., 2018). The aging phenotype in dogs has been previously associated with lower levels of glycine, aspartate, creatine and citrate. Additionally, lower levels of lipoprotein fatty acyl groups were also observed (Richards et al., 2013). There are also metabolic differences with respect to breed, such that large breeds can be separated from others (Viant et al., 2007; Beckmann et al., 2010). Urine seems to more consistently give predictive measures of age, whereas serum analysis yields different metabolites (Richards et al., 2013). Thus far, the only association to weight in metabolomics in the domestic dog has been a negative correlation with tryptophan metabolism (Hoffman et al., 2020).

Urine samples have been shown to be dominated by gut-microbiota metabolites, and serum metabolites may change with respect to the dietary needs of each animal and their feeding state (Lloyd et al., 2016). Of note, the metabolism of primary fibroblasts, which have been shown to retain breed-specific bioenergetic metabolite profiles (Nicholatos et al., 2019), has not been studied from the dual perspectives of aging and breed size. Thus, the aim of this study was use targeted metabolomics, to characterize the metabolome of fibroblasts from young and old, large and small breed dogs.

## Materials and Methods

### Isolation of dog primary fibroblasts

Primary fibroblasts were isolated from puppies and senior dogs of two size classes. The small breed size class included breeds with an adult body mass of 15 kg or less, and the large breed size class included breeds or mixes with an adult body mass of 20 kg or more (Supplementary Table 1). Size classes were based on American Kennel Club (AKC) standards for each breed, as described in Jimenez (2016). Puppy samples were obtained from routine tail docks, ear clips and dewclaw removals performed at veterinarian offices in Central New York and Michigan. Senior dog samples were collected from ear clips immediately after euthanasia. Samples were placed in cold transfer media (Dulbecco’ s modified Eagle medium [DMEM], with 4.5 g/L glucose, 1 mM sodium pyruvate, and 4 mM L-glutamine, supplemented with 10% heat-inactivated fetal bovine serum, and antibiotics [100 U/mL pen/strep], containing 10 mM HEPES) and transferred to Colgate University on ice. To isolate primary fibroblast cells, skin samples were sterilized in 70% ethanol and 10% bleach. Once any fat and bone were removed, skin was minced and incubated in sterile 0.5% Collagenase Type 2 (Worthington Chemicals, Cat. No. LS004176) overnight at 37 °C in an atmosphere of 5% CO_2_ and 5% O_2_. After incubation, the collagenase mixture was filtered through a 20 μm sterile mesh, and centrifuged at 1000 x *g* for 5 min. The resulting supernatant was removed, and the pellet resuspended with 7 mL of mammal media (Dulbecco’ s modified Eagle medium [DMEM], with 4.5 g/L glucose, 1 mM sodium pyruvate, and 4 mM L-glutamine supplemented with 10% heat-inactivated fetal bovine serum, and antibiotics [100 U/mL pen/strep]). Cells were grown in Corning T-25 culture flasks at 37 °C in an atmosphere of 5% O_2_ and 5% CO_2_. When cells reached 90% confluence, they were trypsinized (0.25%) and cryopreserved in liquid N_2_ at 10^6^ cells/mL in DMEM supplemented with 40% fetal bovine serum and 10% dimethylsulfoxide. For experiments, cells were thawed by continuously swirling frozen aliquots in a 37 °C water bath until only a small amount of ice remained, followed by resuspension of each pellet in 6 mL of chilled media, and plating in T-25 flask at 37 °C in an atmosphere of 5% O_2_ and 5% CO_2_.

All the procedures within this study were approved by Colgate University’ s Institutional Care and Use Committee’ s under protocol number 1819-13. Primary fibroblasts from small young (N = 12) and old (N = 9) dogs, and large young (N = 16) and old (N = 12) dogs (see Supplementary Table 1) were expanded from passage 1 (P1) to passage 3 (P3). At P3, cells were trypsinized and pelleted, the supernatant removed and the cells immediately frozen at −80 °C. Frozen cells were shipped on dry ice to the University of Rochester for metabolomics analysis.

### Metabolomics

A targeted LC-MS/MS approach was used (Nadtochiy et al. 2015). Cell pellets were homogenized and metabolites extracted in 3 x 1 mL of ice-cold 80% MeOH. Combined extracts were dried under N_2_ stream and resuspended in 200 μL of 50% MeOH, with 10 µL injected onto HPLC. Protein content of the residual non-extracted material was determined by the Folin Phenol method (Lowry et al. 1957). Reverse phase LC separation utilized a Synergi Fusion-RP column (Phenomenex) and a 3% to 100% methanol ramp over 50 min., on a Prominence 20A HPLC system (Shimadzu). Effluent was directed to a TSQ Quantum Access Max mass spectrometer (Thermo) running a custom library of selected reaction monitoring (SRM) transitions capable of detecting ∼120 common metabolites. Metabolite ID was accomplished by retention time, precursor ion m/z, and at least two product ion m/z’ s obtained at different collision energies. Data was analyzed using Xcalibur software (Thermo).

Samples were analyzed in two batches ∼ 3 mo. apart. The first batch comprised 36 samples run across 3 days and the second batch 22 samples run across 2 days. Pooled daily samples were run before, during and after the samples, as sentinels to monitor instrument sensitivity. One sample was duplicated across both batches, and experimenters were blinded to sample identity until after data analysis was completed. Each batch contained representative samples from each of the 4 groups (small young, small old, large young, large old). Batches were normalized to each other on a per-metabolite basis using average peak size across all groups within a batch. Peak heights were then normalized by the sum of all peaks within each sample, which correlated well with protein content (r^2^ = 0.37, p = 4.6 × 10^−7^). 64 metabolites were reliably identified across 57 samples.

Outliers were identified as those falling outside the 99.99% confidence intervals for each group. As a result, 8 samples were eliminated from further analysis due to excessive noise (>20 outliers among 64 metabolites). Among the remaining 49 samples, discounted outliers and missing values (in total 498 of 3136 possible data points) were imputed on a per-metabolite basis from medians of remaining values within each group (Aittokallio 2011). Data were analyzed using the free web-based *MetaboAnalyst* 4.0 package (Chong et al. 2019). A Benjamini-Hochberg correction is shown in Figure 2 for a false discovery rate (FDR) of 5% (Benjamini and Hochberg 1995).

## Results & Discussion

### Metabolome characteristics

The complete final metabolomic data set for this study (64 metabolites x 49 samples) is available at the data sharing site FigShare (DOI: 10.6084/m9.figshare.14109752). Supplemental Figure 1 shows a pathway analysis plot generated using *Metaboanalyst*, indicating good coverage of several major metabolic pathways and thus permitting conclusions to be drawn about them.

Figure 1 shows a partial least squares difference analysis plot (PLSDA, a dimensionality reduction tool) generated using *Metaboanalyst*, with the top 5 weighted contributors to the principal components shown alongside each axis. Metabolomes of primary fibroblasts from the 4 groups of dogs appeared to cluster into 4 distinct zones, with the sample distributions being similar for each breed size: i.e., small young and small old dogs (yellow and blue) exhibited a horizontal ellipsoid distribution with more variability in component 1 (x-axis), whereas a vertical ellipsoid distribution was seen for large young and large old dogs (green and red) with more variance in component 2 (y-axis). Notably, aging (comparing old vs. young) pushed the distributions in a similar direction (up and to the right) regardless of breed size, similar to the aging patterns in aerobic metabolism seen from primary fibroblast cells of dogs (Jimenez et al., 2018).

**Figure 1.**
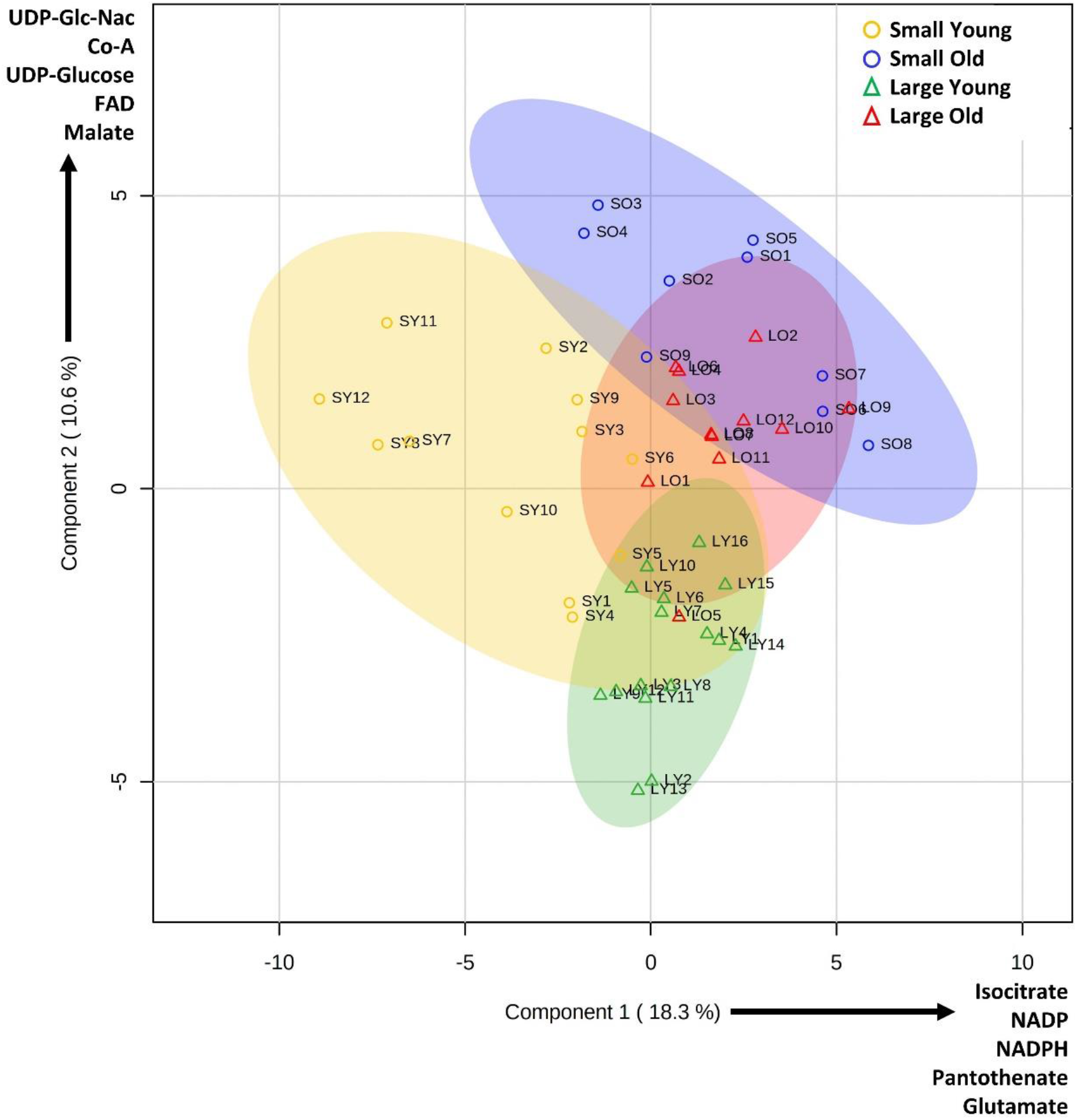
Partial least squares difference analysis (PLSDA) of cells from small, large, young and old dogs. PLSDA was prepares using free *Metaboanalyst* web-based software, incorporating 64 metabolites from 49 samples (see Table 1). Key at upper right denotes symbols for each group. 95% confidence interval areas are indicated by color-appropriate shading. The top 5 metabolite weightings contributing to each principal component are shown alongside the axes.

Figure 2 shows a series of volcano plots, comparing the metabolomes of fibroblasts from the four groups. Panels A and B show comparisons between small and large dogs at either a young (A) or old (B) age. Panels C and D show comparisons between young and old dogs of either a small (C) or large (D) breed size. Overall, both age and size class impacted the metabolome, with some distinct characteristics attributed to each variable. At a young age, there were considerable differences between metabolomes of small vs. large dogs (Figure 2A), whereas far fewer differences were seen between small vs. large in old dogs (Figure 2B). As suggested by the PLSDA plot (Figure 1), this may indicate that aging has a similar impact on the dog metabolome regardless of size class. Nevertheless, some differences were seen between old vs. young dog fibroblasts in both the small (Figure 2C) and large (Figure 2D) breed sizes. These differences are now explored at the metabolic pathway level (below).

**Figure 2.**
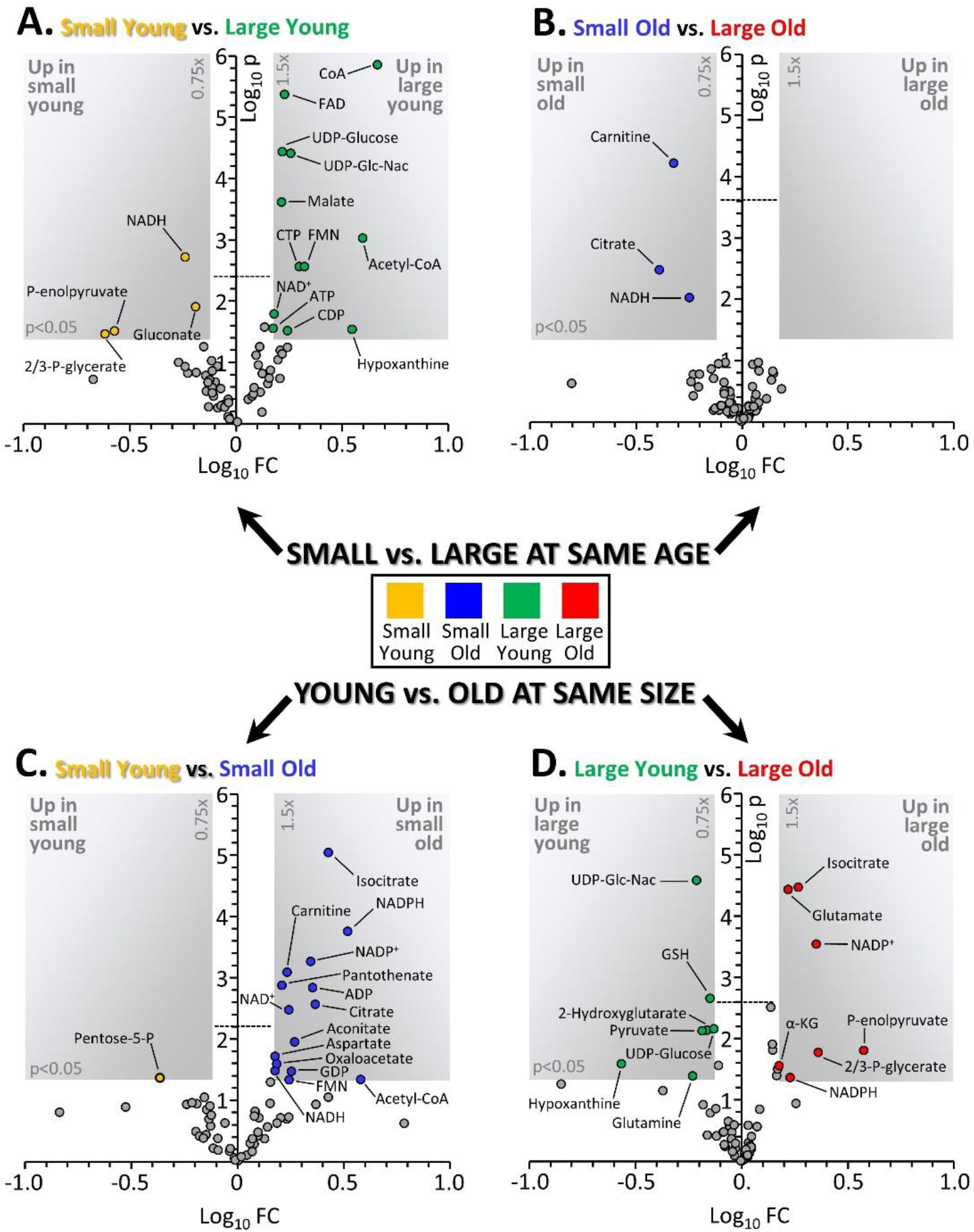
Volcano plots depicting changes in metabolome depending on size or age. Graphs show Log_10_ of the fold change (x-axis) and Log_10_ of the p value from an unpaired heteroscedactic t-test (y-axis) for each metabolite. Gray shaded areas in each plot indicate thresholds of 0.75–1.5 fold change and p=0.05. The horizontal dashed line in each plot indicates the threshold for a Benjamini-Hochberg correction for false discovery rate (Q=0.05). **(A):** Comparison of cells from small young vs. large young dogs. **(B):** Comparison of cells from small old vs. large old dogs. **(C):** Comparison of cells from small young vs. small old dogs. **(D):** Comparison of cells from large young vs. large old dogs. Data points are means, with number of replicates for each group indicated in Table 1. Color coding of groups is as per Figure 1.

### Glycolysis and Associated Pathways

Figure 3 shows glycolysis and its associated metabolic pathways (PPP: pentose phosphate pathway, HBP: hexosamine biosynthetic pathway), with the abundance of those metabolites exhibiting significant differences between groups shown in accompanying graphs.

**Figure 3.**
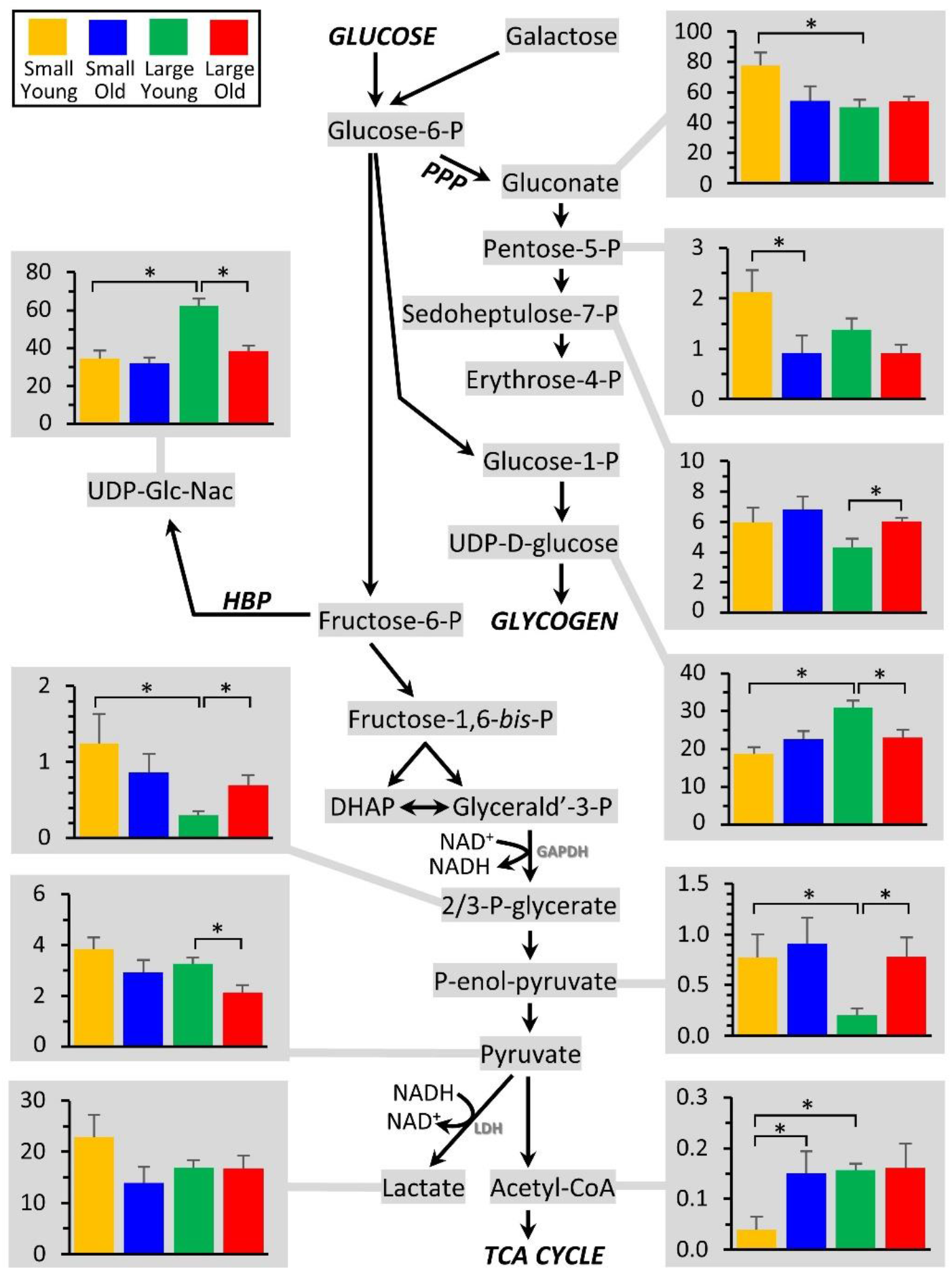
Metabolomics of glycolysis and associated pathways. The glycolytic pathway is shown at center, with all metabolites that were measured highlighted in gray. Those metabolites that exhibited significant differences between groups (Figure 2) are shown as bar graphs, using the same color scheme as Figures 1 & 2. Graphs show means ± SEM of relative metabolite abundance (arbitrary units), with significant differences between groups indicated by asterisks.

At the level of the PPP, several metabolites in the PPP were different between groups, with pentose-5-P being significantly elevated in cells from small young dogs (vs. large young), potentially indicating an altered proliferative potential (given the importance of ribose-P for nucleotide synthesis). In addition, at the level of glycogen metabolism, the precursor UDP-D-glucose was elevated in large young dogs. A similar pattern was observed for UDP-N-acetylglucosamine, the substrate for O-and N-linked glycosylation. Notably all of these differences disappeared with age, consistent with fewer metabolic differences between the small vs. large size classes in old dogs (Figure 2B).

Although no differences were seen in the top half of glycolysis, several notable differences were observed in the lower half of the pathway. In large dogs only, both 2/3-P-glycerate and P-enolpyruvate were elevated in old vs. young dogs, whereas the next metabolite pyruvate was down in old vs. young dogs. To investigate this phenomenon further, a correlation analysis was performed to see if any metabolites correlated with body mass in old animals (N.B. as per Table 1, there was insufficient variation in body mass at the young age group to assess correlations). Indeed, as shown in Supplemental Figure 2, both 2/3-P-glycerate and P-enolpyruvate correlated positively with body mass, whereas the downstream products pyruvate and lactate correlated negatively with body mass, in large old dogs. These data suggest a decrease in pyruvate kinase activity in large old dogs, and are consistent with linear regression analysis of enzymatic activity of this enzyme in blood plasma being lower in large dogs (Wynkoop et al., *in prep*). As such, it may be speculated that the lower half of glycolysis is depressed in larger older dogs, which may represent a metabolic deficiency. Previous work using primary fibroblast cells to measure glycolytic metabolism in small vs. large breeds as they age, found an increase in glycolysis in large breed dogs across their lifespan. This increase could have been due to increases in glucose oxidation or lactate production (Jimenez et al., 2018). Data from the current study supports that increases in glycolysis are associated with glucose oxidation rather than lactate production (since lactate did not change). Using a metabolomics approach, others have found a negative correlation between glycolysis and age in whole-blood samples from dogs (Hoffman et al., 2020)

Further down glycolysis, notably acetyl-CoA was much lower in small young dogs, and pyruvate was also slightly (although not significantly) elevated. While this could indicate a lower activity of pyruvate dehydrogenase in small young dogs, both coenzyme-A and its synthetic precursor pantothenate were also lower in small young dogs (see Figure 7), potentially indicating a depression of coenzyme-A biosynthesis or bioavailability in this group. Nevertheless, such differences appear to resolve in older small dogs, with pantothenate, Co-A and acetyl-CoA all rising with age to levels seen in large dogs.

### TCA Cycle

Figure 4 shows the TCA cycle, with the abundance of those metabolites exhibiting significant differences between groups shown in accompanying graphs. Most of the metabolites in the upper arc of the TCA cycle (malate, OAA, acetyl-CoA, citrate, aconitate, and isocitrate) were all lower in cells from small young dogs, potentially due to low Co-A availability as discussed above. However, such differences were resolved in small old dogs. Since this section of the TCA cycle is also utilized to export citrate from mitochondria to make fatty acids (via ATP citrate lyase), this may also indicate a greater draw on the cycle for biosynthetic purposes (and consistent with the elevated pentose-5-P in the PPP, see Figure 3). Others have found a negative correlation between the TCA cycle components and age using whole-blood samples from dogs (Hoffman et al., 2020)

**Figure 4.**
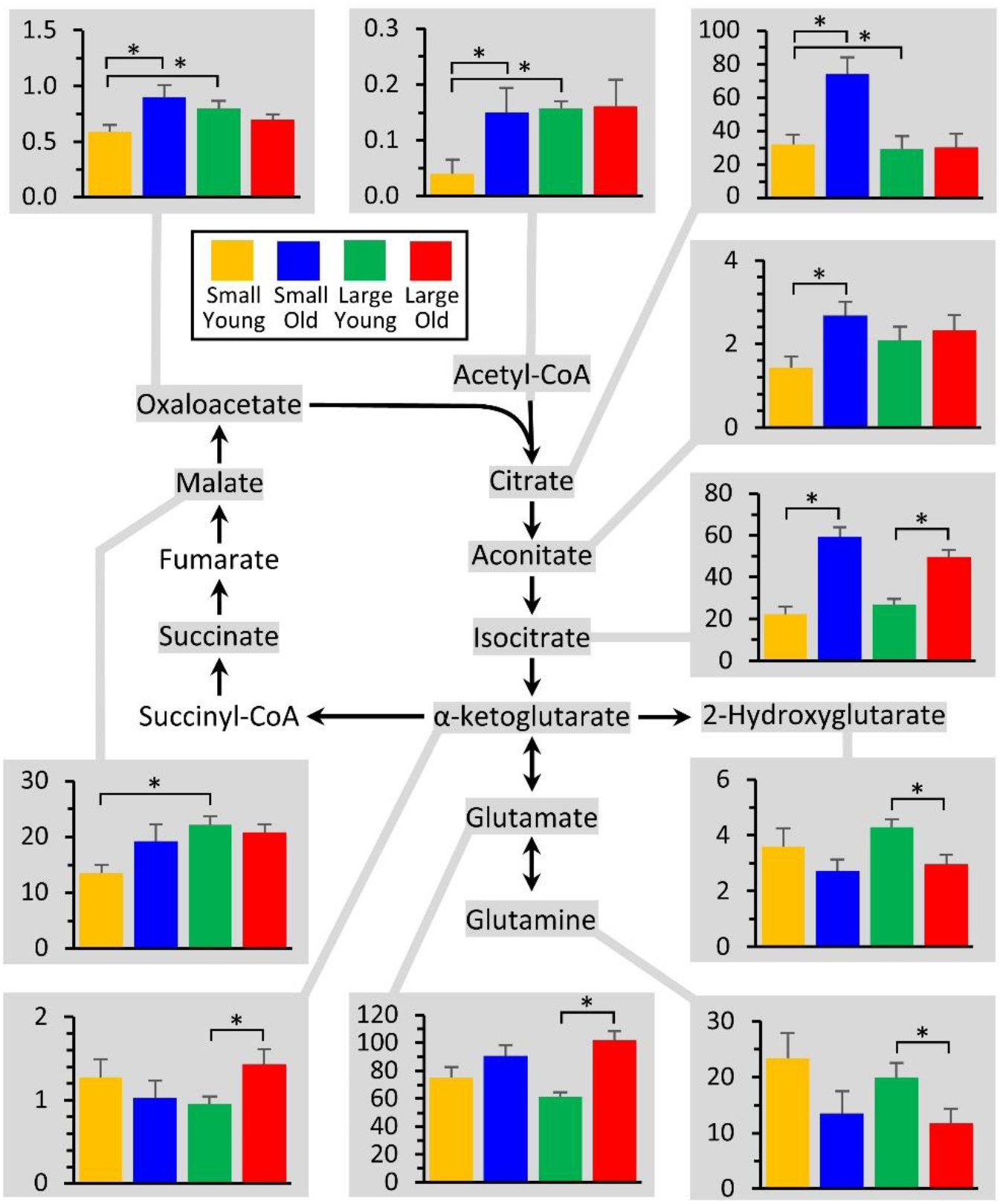
Metabolomics of the TCA cycle. The TCA cycle is shown at center, with all metabolites that were measured highlighted in gray. Those metabolites that exhibited significant differences between groups (Figure 2) are shown as bar graphs, using the same color scheme as other Figures. Graphs show means ± SEM of relative metabolite abundance (arbitrary units), with significant differences between groups indicated by asterisks.

For the lower arc of the TCA cycle, notably old large dogs exhibited lower levels of glutamine, and elevated levels of glutamate and α-ketoglutarate. If indeed large old dogs are deficient in the lower half of glycolysis (see above), this may indicate a greater degree of anaplerosis from glutamine, to ensure sufficient carbon flux into the TCA cycle.

Notably the *oncometabolite* 2-hydroxyglutarate (2-HG) was lower in both small and large old dogs. Although this metabolite has been proposed as a biomarker for certain types of cancer, this applies only to the D isomer of 2-HG (i.e. D-2-HG) which originates from mutated forms of isocitrate dehydrogenase (Ward et al., 2010). By contrast, L-2-HG can be made by several metabolic enzymes under conditions of hypoxia or metabolic acidosis (Oldham et al., 2015; Nadtochiy et al., 2016). Unfortunately, the methods used herein were incapable of distinguishing L-vs. D-2-HG, so we are unable to tie this metabolic difference to any differences in cancer phenotype as dogs age.

### Malate-Aspartate Shuttle

The glycolytic enzyme glyceraldehyde-3-phosphate dehydrogenase (GAPDH) reduces NAD^+^ to NADH, and the reoxidation of this NADH back to NAD^+^ is a critical determinant of glycolytic flux (Figure 3). NADH is usually reoxidized via two mechanisms: either lactate dehydrogenase in the cytosol (LDH, Figure 3), or by transferring the reducing equivalency of NADH into mitochondria via malate-aspartate shuttle (MAS, Figure 5). A deficiency in pyruvate kinase in large old dogs (see above) could render the LDH reoxidation pathway less available, which may drive a greater reliance on the MAS to reoxidize glycolytic NADH. Figure 5 shows that several components of the MAS were altered both by age and breed size. However, since these studies were conducted on whole cell extracts, it is not possible to determine whether the metabolites measured herein originated from the cytosolic or mitochondrial compartment. As such, further investigation is required to determine the status of the MAS in aging dogs.

**Figure 5.**
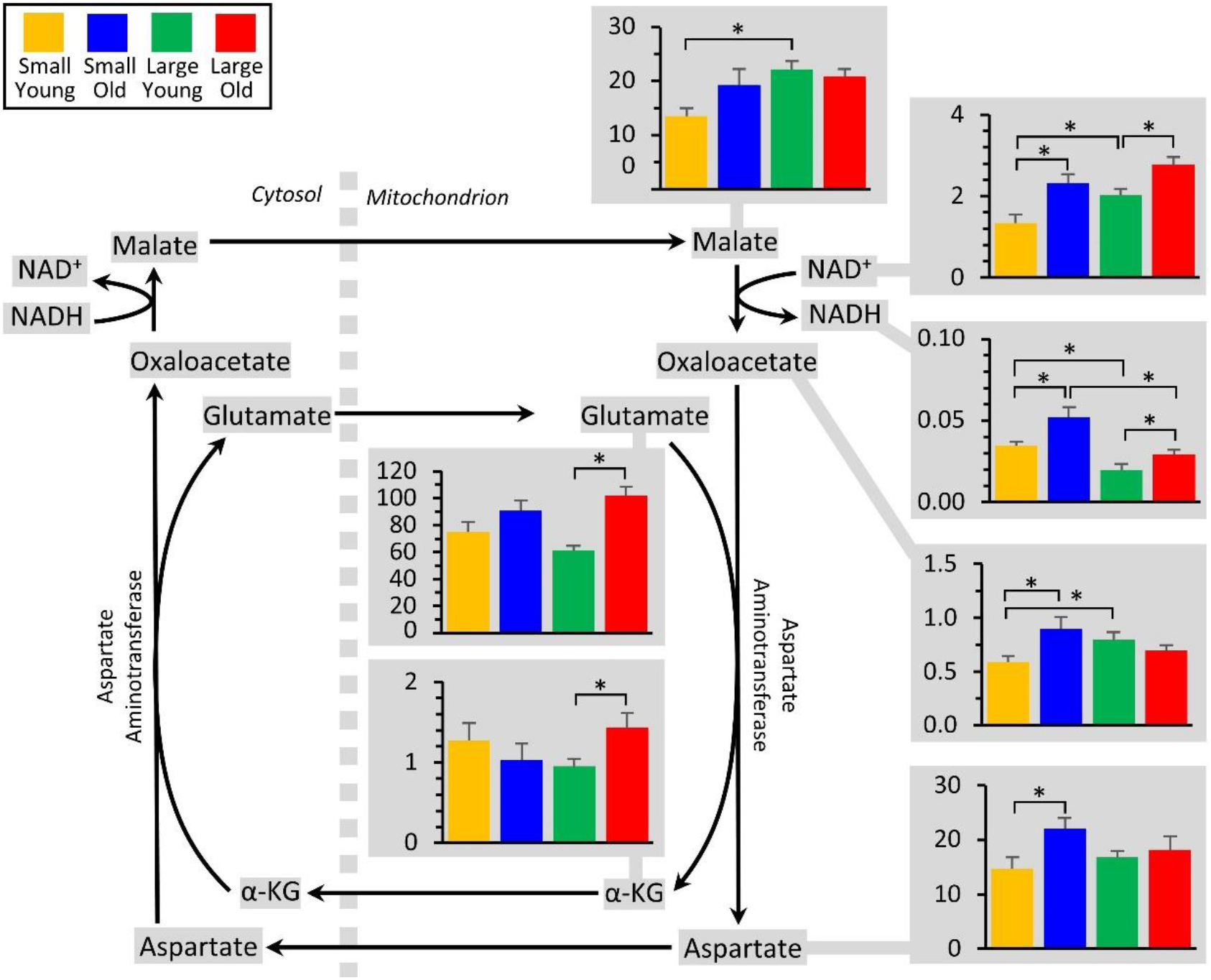
Metabolomics of the malate/aspartate shuttle. The shuttle is shown at center, with all metabolites that were measured highlighted in gray. Those metabolites that exhibited significant differences between groups (Figure 2) are shown as bar graphs, using the same color scheme as other Figures. Graphs show means ± SEM of relative metabolite abundance (arbitrary units), with significant differences between groups indicated by asterisks. Note that several metabolites in the shuttle are present in both mitochondrial and cytosolic compartments, and the methods employed herein were incapable of distinguishing between localized metabolite pools.

### Redox Metabolites

Oxidative stress is believed to be an important (although somewhat controversial) component of the aging process (Lee et al., 2014). Consistent with a greater degree of oxidative stress in aging animals, Figure 6 shows that cells from both small and large breeds exhibited lower levels of the antioxidant reduced glutathione (GSH) in old vs. young, which contrasts with findings in Jimenez et al. (2018). These two studies include differing sample sizes and differing methods of measuring GSH, which may lead to the discrepancy. Using a metabolomics approach, others have found that GSH in plasma from domestic dogs was lower in smaller dogs compared with larger dogs (Middleton et al., 2017), and there was a negative correlation with age and GSH using whole-blood (Hoffman et al., 2020). No changes were seen in oxidized glutathione (GSSG), such that glutathione redox state (GSH/GSSG) also declined with age in both breed size groups (*see important note below).

**Figure 6.**
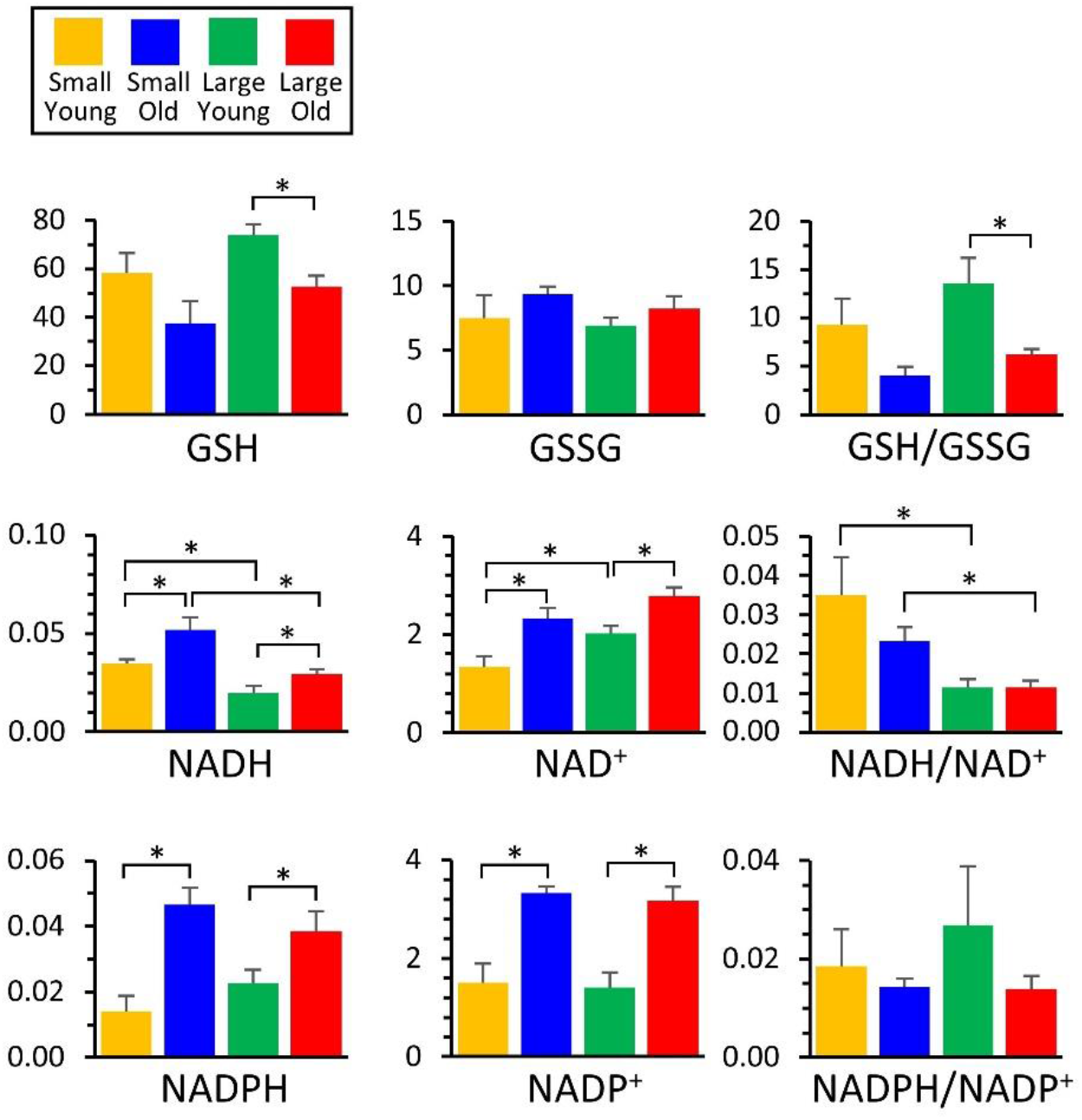
Metabolomics of redox. Those metabolites that exhibited significant differences between groups (Figure 2) are shown as bar graphs, using the same color scheme as other Figures. Graphs show means ± SEM of relative metabolite abundance (arbitrary units), with significant differences between groups indicated by asterisks. Note that although redox ratios (e.g., GSH/GSSG) are shown, these are for illustrative purposes only, since different electrospray ionization efficiencies for each metabolite render such ratios non-quantitative.

Pyrimidine nucleotides also exhibited some interesting differences between the 4 groups. Both NADH and NAD^+^ levels appeared to increase with age in small and large dogs. While this would appear to be inconsistent with recent reports that a decline in bulk NAD^+^ levels is an important component of the aging process (Zhang et al., 2016), it is notable that these increases were generally of lower magnitude in the large dogs, compared to small dogs, which is consistent with a shorter lifespan in large dogs. Similar increases in nucleotide abundance with age were seen for NADPH and NADP^+^, and again these increases were of greater magnitude in smaller dogs.

*Although redox ratios of GSH/GSSG, NADH/NAD^+^ and NADPH/NADP^+^ are shown in Figure 6, the nature of the LC-MS/MS based measurement system is such that quantitation is relative between groups or samples for the same metabolite. However, differences in ionization properties do not permit comparisons to be made between metabolites. As such, these ratios are shown for informational purposes only and no conclusions should be drawn from them.

### Bioenergetics and co-factors

As shown in Figure 7, Both ATP and ADP appeared to increase with age in small dogs, but their levels were already high in large dogs at a young age. As expected for a general decline in bioenergetic function with age, the ATP/ADP ratio was lower in old dogs from both breed size groups (see caveat above regarding metabolite ratios). Similar patterns in bulk nucleotide levels were seen for guanosine, cytosine and uridine nucleotide phosphates (not shown in figures, see full data set for details), all of which became elevated with age in small dogs, but were already elevated in old dogs. Given the role of such nucleotides in proliferation and DNA/RNA replication, these findings may reflect a lower proliferative potential in small young dogs. It is interesting to juxtapose these findings with that of elevated pentose-5-P levels in small young dogs (see above), perhaps suggesting a defect in nucleotide biosynthesis (more substrate, less product).

**Figure 7.**
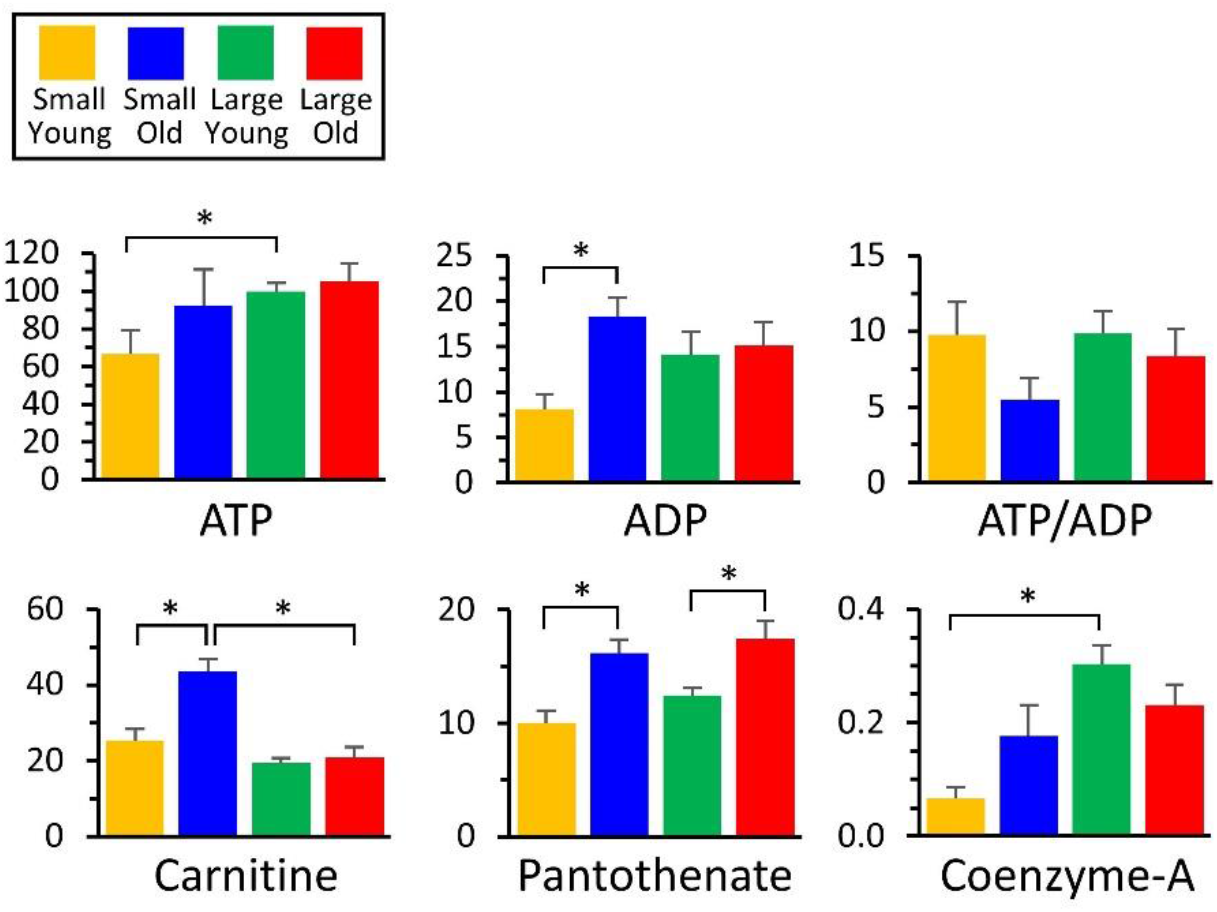
Metabolomics of purine nucleotides and co-factors. Those metabolites that exhibited significant differences between groups (Figure 2) are shown as bar graphs, using the same color scheme as other Figures. Graphs show means ± SEM of relative metabolite abundance (arbitrary units), with significant differences between groups indicated by asterisks. Note that although ratios (e.g., ATP/ADP) are shown, these are for illustrative purposes only, since different electrospray ionization efficiencies for each metabolite render such ratios non-quantitative.

Another notable finding was a significant elevation in carnitine as small dogs aged, which was not seen in large dogs. This could reflect the bioavailability of carnitine in the diet (e.g., red meat is rich in carnitine), and indeed Middleton et al. (2017) reported enhanced protein digestibility in small dogs. Given the role of carnitine in the import of fatty acids to mitochondria for β-oxidation, a failure to elevate carnitine in old large dogs could lead to a depression in fat oxidation. This may partly underlie the apparent upregulation of anaplerosis into the TCA cycle seen in large old dogs (Figure 4). Using metabolomics in primary fibroblast cells of different dog breeds, Nicholatos et al., (2019) found that longer-lived (smaller) breeds have lower concentrations of acylcanitines, and higher fatty acid oxidation, which would agree with our current findings. L-carnitine is also a metabolite relevant to aging in humans (Lee et al., 2014). However, most of the human aging literature suggests a correlation between lipid metabolism and aging (Lee et al., 2014), which was not seen in metabolomics of primary fibroblast cells from dogs. Additionally, Hoffman et al., (2020) demonstrated a positive correlation between saturated fatty acid β-oxidation and age in whole blood samples of dogs.

Finally, as noted above (glycolysis), age-dependent increases in the co-enzyme A precursor pantothenate were seen in both breed sizes, but coenzyme-A itself did not increase in large old dogs. This could indicate a defect on Co-A biosynthesis, although it does not appear to impact the levels of acetyl-CoA originating from upstream metabolic pathways (glycolysis or the oxidation of fat, ketones or branched-chain amino acids).

### Summary

Herein, several significant differences were observed between the metabolic profiles of fibroblasts obtained from young and old, large and small dogs. In general, regardless of size, older dogs exhibited a lower glutathione redox state (GSH/GSSG) and lower bioenergetic state (ATP/ADP), consistent with numerous previous reports on the metabolic consequences of the aging process. In addition, several size-specific metabolic patterns were observed, with divergence between small and large dogs with aging. This included: (i) An apparent defect in the lower half of glycolysis in large old dogs, at the level of pyruvate kinase. (ii) Increased glutamine anaplerosis into the TCA cycle in large old dogs. (iii) A potential defect in co-enzyme A biosynthesis in large old dogs. (iv) Age-dependent increases in nucleotides and nucleotide phosphates in small dogs, bringing their levels to those already seen in large dogs at all ages. (v) An age dependent increase in carnitine in small dogs that was absent in large dogs. These differences are summarized in Figure 8. The underlying enzymatic origins of these observed differences in steady-state metabolomes are the subject of ongoing investigations, aimed at revealing potential therapeutic avenues to impact the aging process in domestic dogs.

**Figure 8:**
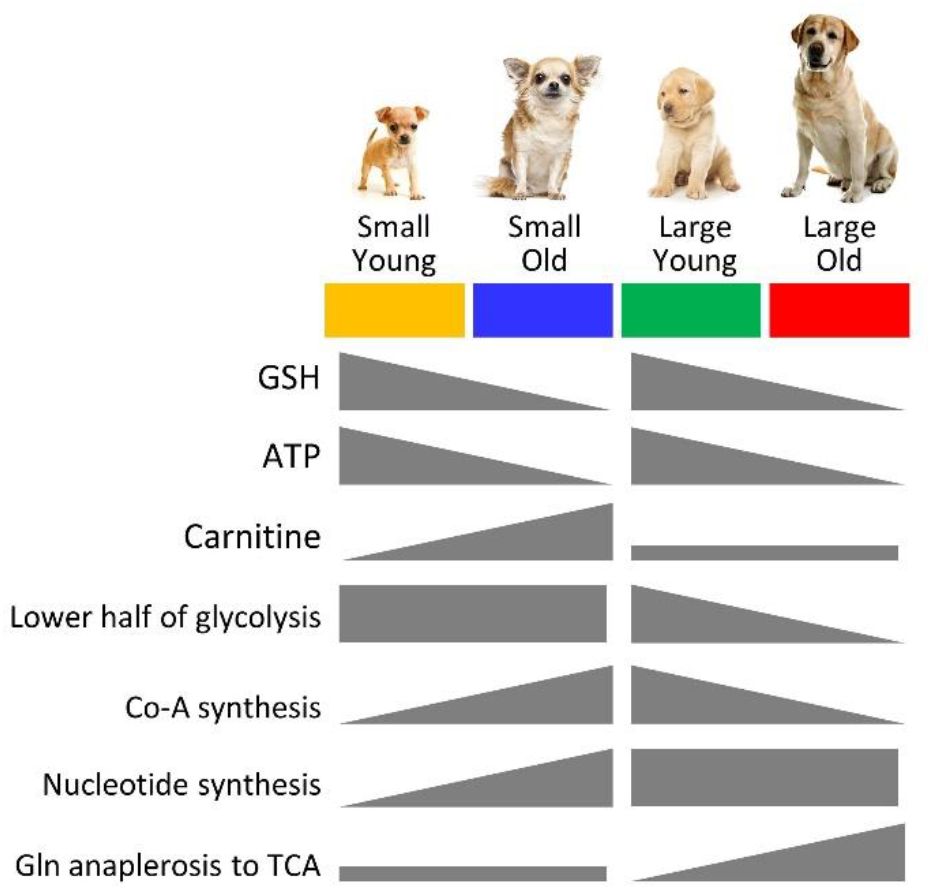
Summary of metabolomic changes with age in small and large dogs. Using the color scheme from other figures, the main changes in metabolites or key metabolic pathways are shown as increasing or decreasing, in the gray symbols below.

## Acknowledgements

We are grateful to the following veterinarians and veterinary practices for providing us with samples: Dr. Kerri Hudson, Dr. James Gilchrist, Dr. Heather Culbertson and Morgan Peppenelli at Waterville Veterinary Clinic (New York); Dr. Frank Capella from Village Vet in Wampsville, NY. Pet Street Station Animal Hospital (New York); Dr. Jim Bader at Mapleview Animal Hospital (Michigan). We are also grateful to the following breeders for participating in our study: Rhonda Poe, Bob Stauffer, Allison Mitchell, Nancy Secrist, Valeria Rickard, Joanne Manning, Lita Long, Betsy Geertson, Susan Banovic, Lisa Uhrich, Sheryl Beitch, Al Farrier, Barbara Hoopes and Rachel Sann. Work in the lab of PSB is funded by a grant from the US National Institutes of Health (R01-HL071158).

