## Supplemental Information for "Metabolomics of aging in primary fibroblasts from small and large breed dogs"

1 Table

2 Figures

| Breed | Age | Size class | Age class | N |
|---|---|---|---|---|
| Alaskan Malamute mix | 15 y/o | Large | Old | 1 |
| Boxer | 13.5 y/o | Large | Old | 1 |
| Boxer/Sheperd | 13 y/o | Large | Old | 1 |
| Golden retriever | 11 and 13 y/o | Large | Old | 2 |
| Labrador mix | 14 y/o | Large | Old | 1 |
| Pibull | 15 y/o | Large | Old | 1 |
| Australian Shepherd | 13 and 15.5 y/o | Large | Old | 2 |
| Rottie/Shepherd | 10 y/o | Large | Old | 1 |
| Siberian Husky | 6 and 11 y/o | Large | Old | 2 |
| Airdale terrier | 5 days old | Large | Young | 2 |
| Australian cattle dog | 3 days old | Large | Young | 2 |
| Doberman | 9 weeks old | Large | Young | 1 |
| German short-haired pointer | 2 days old | Large | Young | 2 |
| German wire-haired pointer | 2 days old | Large | Young | 2 |
| Labradoodle | 3 days old | Large | Young | 2 |
| Old English sheepdog | 3 days old | Large | Young | 3 |
| Rottweiler | 2 days old | Large | Young | 1 |
| Standard poodle | 4 days old | Large | Young | 1 |
| Bichon Frise | 12 y/o | Small | Old | 1 |
| Cocker Spaniel mix | 14 y/o | Small | Old | 1 |
| Jack Russell Terrier | 14 y/o | Small | Old | 1 |
| Shetland sheepdog | 14, 14, and 12 y/o | Small | Old | 3 |
| Shih Tzu | 14 y/o | Small | Old | 1 |
| Shih Tzu mix | 13 y/o | Small | Old | 1 |
| Yorkshire terrier | 17 y/o | Small | Old | 1 |
| Cavalier King Charles Spaniel | 4 days old | Small | Young | 2 |
| Corgi | 1 day old | Small | Young | 2 |
| Havenese | 4 days old | Small | Young | 2 |
| Soft coated wheaten terrier | 3 days old | Small | Young | 2 |
| Toy poodle | 3 days old | Small | Young | 2 |
| Yorkshire terrier | 3 days old | Small | Young | 2 |

**Table 1.** Information about breeds, ages, size and age classes, and sample sizes for dogs included in this study.

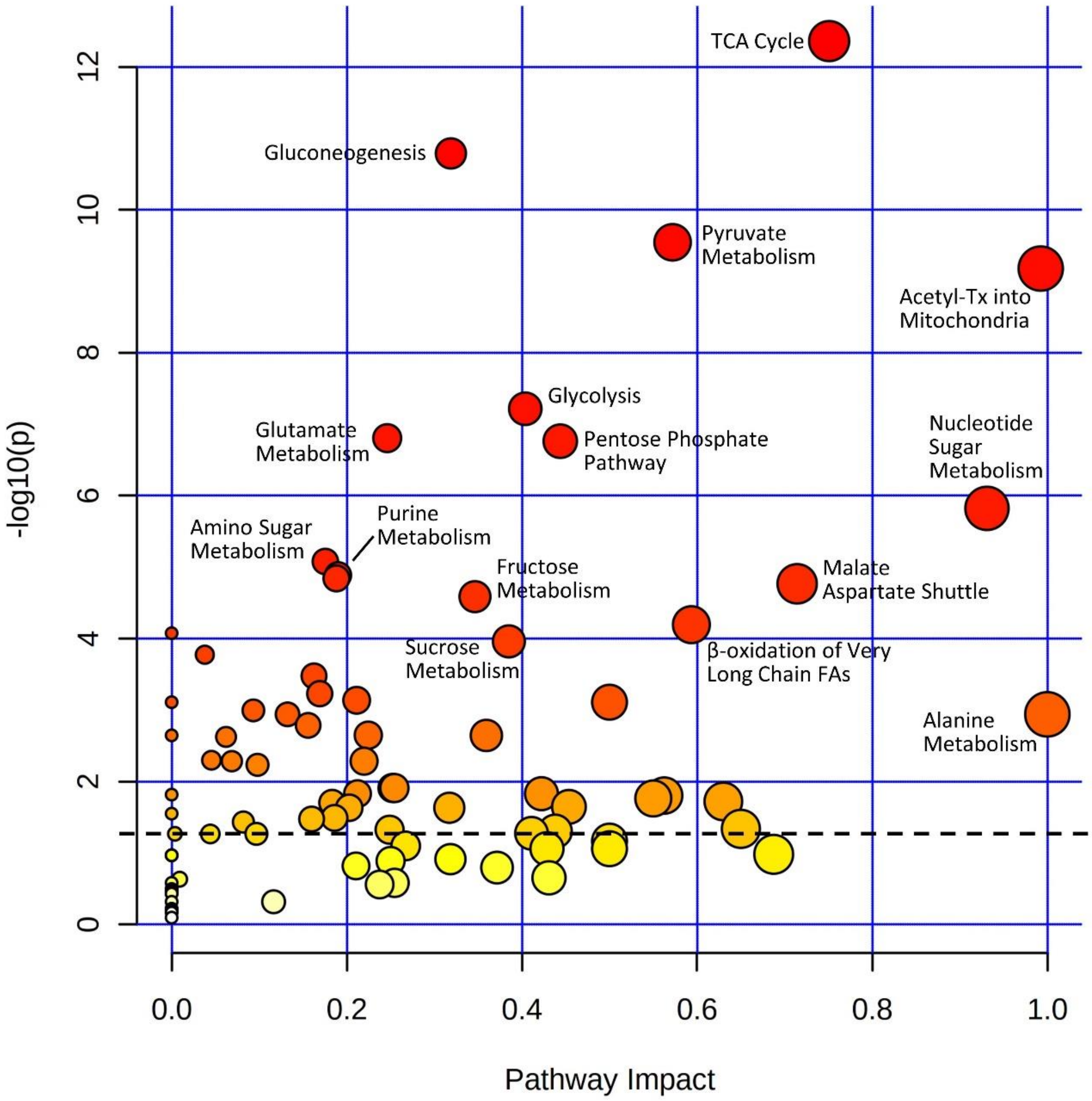


**Supplemental Figure 1. Pathway analysis for the metabolomic data set.** Pathway Analysis was prepared using free web-based Metaboanalyst software. Y-axis shows -Log$_{10}$ of p-values from the pathway enrichment analysis (Log10 of p=0.05 is 1.3, so pathways above the dashed line were considered to be significantly enriched in the analysis). X-axis shows pathway impact values from pathway topology analysis, indicating the degree of contiguous coverage within a pathway.

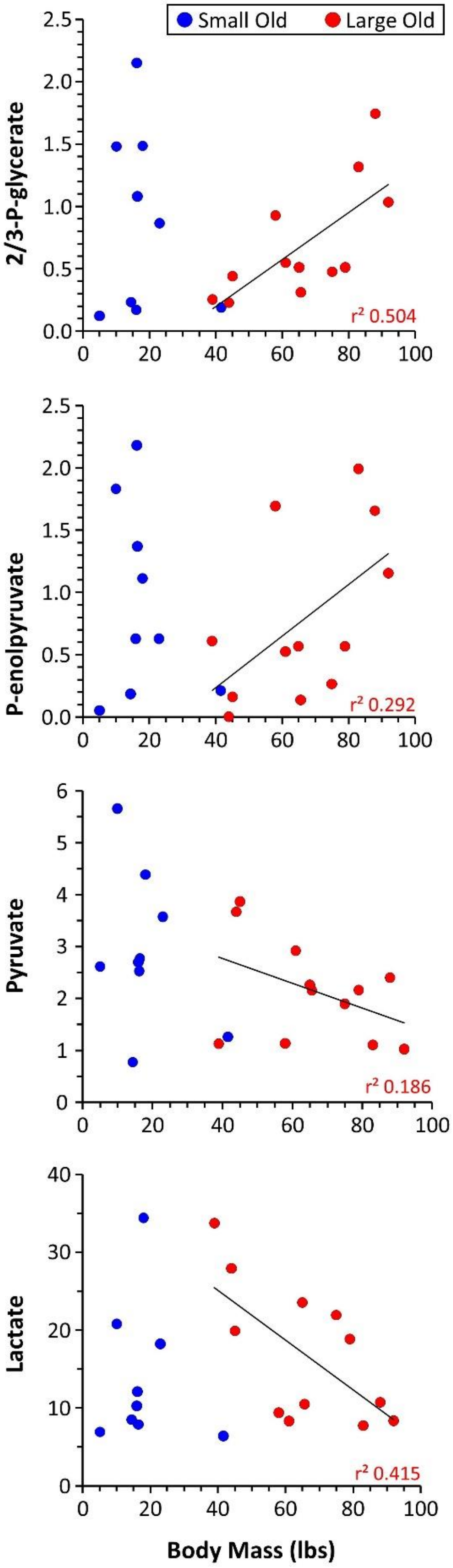


**Supplementary Figure 2. Correlation of metabolites with body mass in old dogs.** Correlations are shown between body mass and relative abundance (arbitrary units) of four glycolytic metabolites, in small and large old dogs. Due to insufficient variation in body mass, correlations were not calculable in young dogs of either breed size. Linear fit trendline and square of Pearson product moment correlation coefficient ($r^2$) for the large old dog data set are shown on each graph.
